# A molecular network of the aging brain implicates *INPPL1* and *PLXNB1* in Alzheimer’s disease

**DOI:** 10.1101/205807

**Authors:** S. Mostafavi, C. Gaiteri, S. E. Sullivan, C.C. White, S. Tasaki, J. Xu, M. Taga, H. Klein, E. Patrick, V. Komashko, C. McCabe, R. Smith, E.B. Bradshaw, D. Root, A. Regev, L. Yu, L.B. Chibnik, J.A. Schneider, T. Young-Pearse, D.A. Bennett, P.L. De Jager

**Author notes:** Contributed equally (S. Mostafavi, C. Gaiteri, D.A. Bennett, P.L. De Jager).

## Abstract

The fact that only symptomatic therapies of small effect are available for Alzheimer’s disease (AD) today highlights the need for new therapeutic targets with which to prevent a major contributor to aging-related cognitive decline. Here, we report the construction and validation of a molecular network of the aging human frontal cortex. Using RNA sequence data from 478 individuals, we first identify the role of modules of coexpressed genes, and then confirm them in independent AD datasets. Then, we prioritize influential genes in AD-related modules and test our predictions in human model systems. We functionally validate two putative regulator genes in human astrocytes: *INPPL1* and *PLXNB1*, whose activity in AD may be related to semaphorin signalling and type II diabetes, which have both been implicated in AD. This arc of network identification followed by statistical and experimental validation provides specific new targets for therapeutic development and illustrates a network approach to a complex disease.

**One sentence summary:** Molecular network analysis of RNA sequencing data from the aging human cortex identifies new Alzheimer’s and cognitive decline genes.

## Introduction

The incidence of late-onset Alzheimer’s disease (AD) is expected to triple by 2050^1^, yet no therapies are available to treat or prevent the disease^2^. Possible reasons for the continued failure of AD trials include the biological complexity of the disease coupled with its phenotypic heterogeneity^3^. Recent genome-wide association studies (GWAS) have identified new potential therapeutic targets involved in endocytosis, metabolism, and inflammation^4^. However, it has been difficult to transition from mostly non-coding genetic variants that influence AD risk to molecular and cellular mechanisms that lead to the characteristic accumulation of β-amyloid and helical filament tau (PHFtau) pathology as well as the subsequent cognitive decline of AD. Furthermore, as large GWAS generally do not distinguish between variants related to AD neuropathology versus unknown contributors to cognitive decline, additional molecular, pathological and cognitive measures are likely necessary to determine the drivers of various aspects of AD.

Here, we describe an analysis of participants from two large longitudinal cohort studies of aging (total n=478), as well as a validation cohort (n=82), which have careful assessments of both antemortem cognitive function and postmortem neuropathologic burden. We hypothesized that using RNA-sequencing (RNA-Seq) data from the dorsal lateral frontal cortex (DLPFC) enabled us to identify coherent intermediate cellular mechanisms that are associated with cognitive decline and/or neuropathological changes. Noting that dynamic genomic measurements such as RNA-Seq reflect the collective effect of upstream, downstream, and disease-correlated processes, we use a series of network-based approaches that account for known and hidden confounding factors in order to enrich our results for likely upstream associations.

As cellular functions are carried out by networks of interacting molecules, uncovering these network's structures can provide clues about cellular functions^5,6^. Such a network-based perspective also provides a more nuanced molecular definition of complex disease than do traditional single-gene associations^7–9^ because it provides a natural framework to assemble disparate single gene findings into disease mechanisms^10–12^. Transcriptomic data can be used to build such networks, facilitating the identification of groups of coexpressed genes or “modules” that represent cellular processes and can be related to phenotypes of interest^13,14^. Compared to traditional single gene or post-hoc pathway enrichment approaches, coexpression network-based analysis offers an unsupervised and tissue-specific approach that identifies cellular subnetworks related to disease phenotypes, independent of historical bias arising from research on particular genes and pathways^14^.

Coexpression approaches have been applied to AD and have identified genes associated with a syndromic diagnosis of AD dementia^*15,16*^ However these approaches have never been applied to large cohorts with quantitative measurements of both AD, other common neuropathologies, and cognitive decline. Because neuropathology and cognition show important divergence in AD^17,18^, jointly modeling both aspects of the disease may better capture the relationship between molecular events and different stages of the disease process. Further, previous efforts did not distinguish between transcriptomic patterns that are indirectly associated with AD phenotypes, via a chain of intermediaries, from those that are directly associated^15^. Here, we build on prior work to address these limitations. Our approach, called gene <u>m</u>odule <u>t</u>rait <u>n</u>etwork analysis (MTN) (**Figure 1**), constructs gene expression modules and identifies those that are directly associated with cognitive decline, conditioned on neuropathology and other large-scale transcriptomic changes in the aging brain. We confirm the biological plausibility of this systems biology analysis in five other types of independent datasets. Finally, we test the identified associations in a relevant human model system to functionally characterize selected genes and prioritize candidates for further drug development.

**Fig 1.**
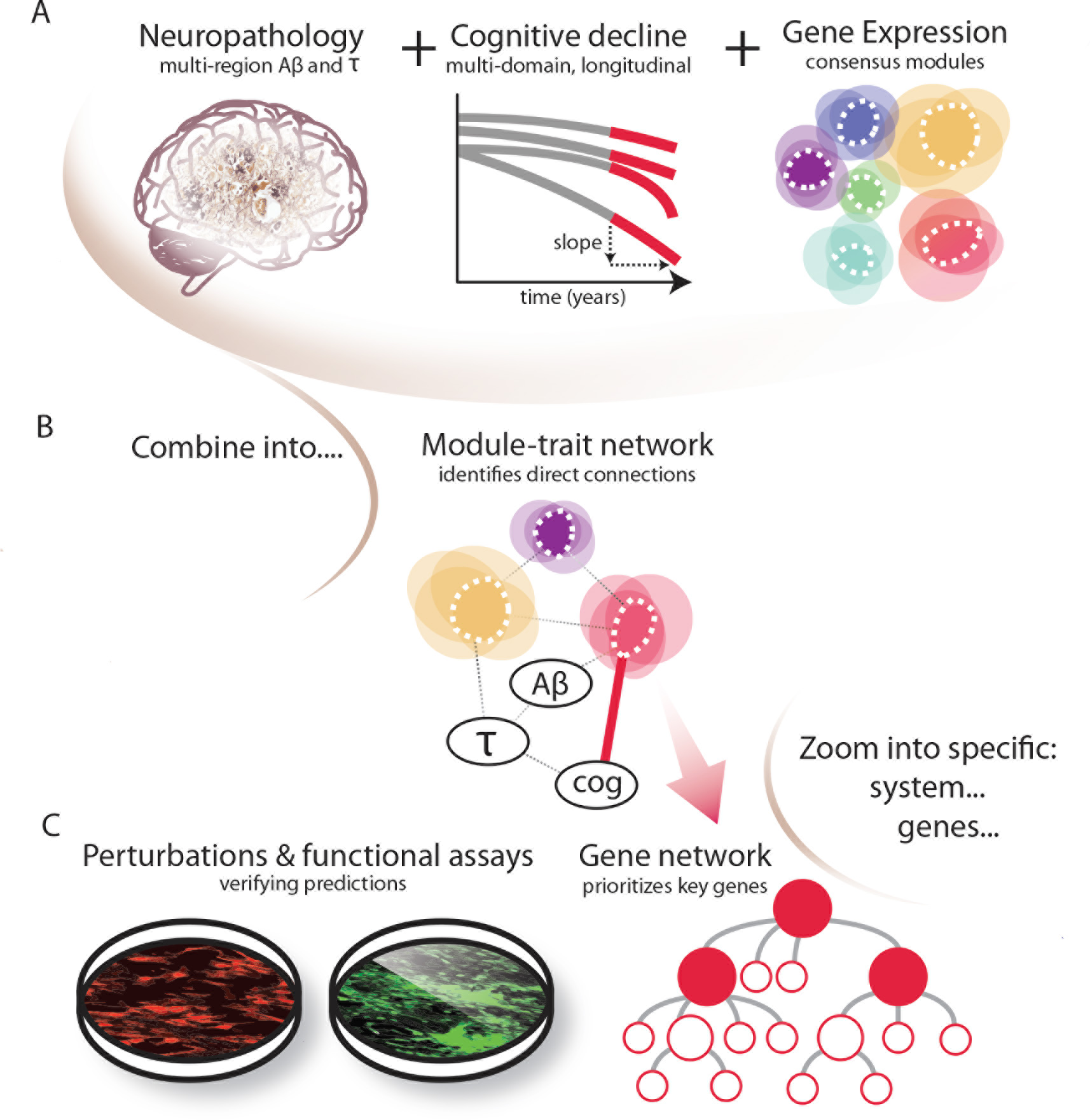

## Results

### Data origin and phenotypes

Data were derived from subjects enrolled in the Religious Orders Study (ROS) or the Rush Memory and Aging Project (MAP), two prospective clinical-pathologic cohort studies of aging and dementia. All participants are non-demented at enrolment, have annual detailed clinical evaluations and have agreed to brain donation. At death, each brain undergoes a structured, quantitative neuropathologic assessment (see **Supplementary methods**). The two studies (collectively referred to as ROSMAP) share clinical and neuropathological standards, allowing for joint analyses of the data. Individual trajectories of cognitive decline are calculated from longitudinal cognitive measures that include up to 20 yearly evaluations^19^. For this study, we used data from 478 participants, of which 257 (62.5%) are women. The mean age at death was 88.7 years (95% confidence interval 88.1-89.3). Over the course of the study, some subjects experience cognitive decline, and, at the time of death, 32% remain cognitively nonimpaired, 27% have mild cognitive impairment (MCI), 39% have a diagnosis of AD dementia (with or without a co-morbid condition), and 2% have another form of dementia. When brain pathology is measured in all of these persons, 58% of subjects receive a diagnosis of pathologic AD. However, of these individuals with pathologic AD, 46% are clinically non-demented (they are either cognitively non-impaired or have MCI), illustrating the well-described divergence of pathologic and clinical diagnoses of AD dementia (**Table S1**).

Our analysis includes five phenotypic traits related to AD: two of these traits are clinical measures - a clinical diagnosis of AD dementia proximate to death (ClinAD), and a continuous measure of cognitive decline over time quantified as a per-subject slope of the cognitive decline trajectory from a linear mixed-effects model^20^ (“cognitive decline” abbreviated as “Cog Dec”). The three pathology variables include continuous measures of PHFtau tangle density and β-amyloid burden (both averaged over multiple regions) and a binary diagnosis of pathologic AD (PathoAD) (See the **Supplementary methods** for details).

In these subjects, we see the expected strong association of *APOE ɛ4* with ClinAD (p=5.55x10^−16^; used logistic regression to model the number of *ɛ4* while accounting for age and sex), but even this unique genetic risk factor explains little of the variance in ClinAD (2.2% variance explained) or cognitive decline (5.1% variance explained). Subsequent inclusion of the other 21 AD susceptibility variants to represent the known genetic architecture of AD only explains 2.1% of ClinAD and 7.6% of cognitive decline. Thus, while these robustly validated susceptibility variants provide important insights into risk factors that contribute to AD, they capture only a small fraction of the biology of the disease, much of which may be influenced by non-genetic risk factors. The transcriptome, with its dynamic nature that is molded by environmental exposures and life experiences, provides a complementary approach to therapeutic target discovery in AD.

### RNA-Seq gene expression data and standard association analysis

After rigorous quality control evaluations, we retained RNA-Seq data from the dorsolateral prefrontal cortex (DLPFC) of 478 individuals for downstream analyses; an average of 95 million (median 90 million) paired-end reads was available for each subject. RNA-Seq data were then normalized to account for the effect of a large number of known biological and technical confounding factors. Genes with low expression were removed to reduce the influence of technical noise, resulting in quantified expression for 13,484 unique genes (25,400 transcripts) (**Supplementary methods**).

We first performed a standard transcriptome-wide association study (TWAS) to identify genes whose expression levels associate with the clinical and pathologic AD-related phenotypes. We found that the expression levels of thousands of genes associate (FDR<0.05) with one or more of these traits (**Supplementary Table S2; Figure 2A**). The combination of cognitive and pathological phenotypes allows us to compare their respective effects on gene expression, and we observed that cognitive decline was associated with the largest number of genes (3,025 genes at FDR<0.05), compared to the other AD traits. Accordingly, the key clinical variable of cognitive decline may implicate additional molecular mechanisms in AD, beyond those found by neuropathological markers of AD. Indeed a majority of age-related cognitive decline cannot be accounted for by current measures of AD or other age-related pathologies^21^. Overall, using the π_1_ statistic^22^, we estimated that 55-90% of the associated genes are shared amongst these correlated AD traits (**Figure S2A**). The large number of associated genes highlights the need to assemble these finding into coherent biological processes that are directly associated with specific disease-relevant endpoints.

**Fig 2.**
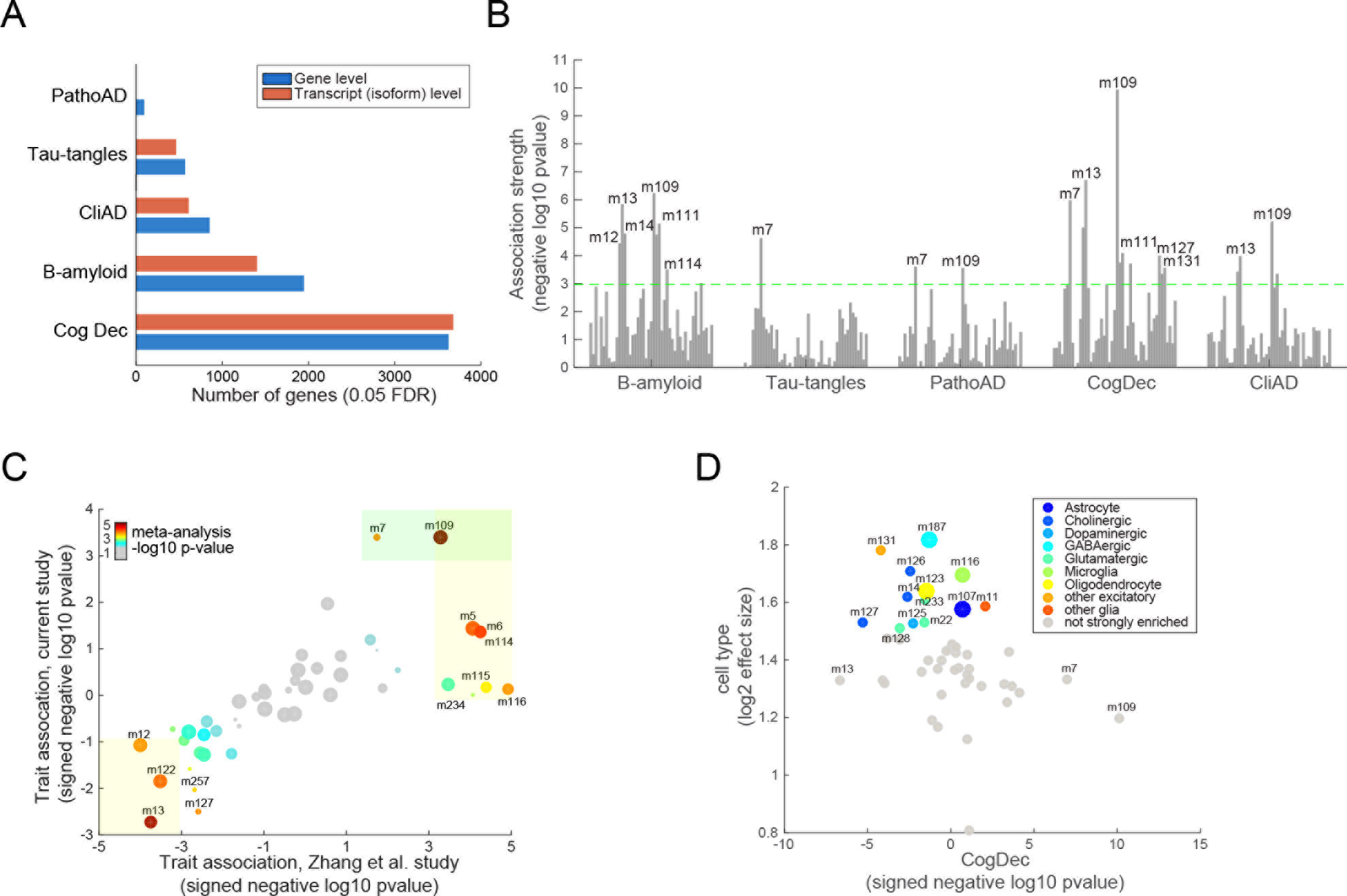

### Constructing the nodes of our module-trait network

In order to identify coherent cellular processes that impact AD phenotypes, we utilize a module-trait network (MTN) approach. The goal of this approach is to go beyond single gene level associations in defining robust molecular mechanisms while avoiding the limitations of pathways derived from ontology databases. MTN summarizes large-scale transcriptome changes into gene modules (or clusters) and prioritizes specific genes within a module for additional experiments. MTN consists of three steps. As described in detail below, we first identify groups of coexpressed genes or “modules”, which are validated in other datasets. Coexpressed gene sets represent the outcome of transcriptional regulatory mechanisms that include transcription factors, chromatin conformation and other effects; coexpression can also stem from the presence of latent factors that generate correlations among groups of genes, such as the proportion of different cell types present in the sampled tissue^14^. In the second step, we identify which modules have direct relationships with cognitive decline and other AD traits using Bayesian networks to prune correlations between modules and AD-related traits that are indirect. In the final analytical step, we select a top-scoring module and prioritize genes that are uniquely influential *within* that module for validation in our *in vitro* model systems.

In the first step of MTN, we applied the SpeakEasy consensus clustering algorithm^23^ to our RNA-Seq data and identified 47 mutually exclusive modules ranging in size between 20 and 556 gene members (**Table S3; Figure S4; Supplementary methods**). We note that the clustering assignment identified by the SpeakEasy algorithm overlapped significantly with those proposed by the frequently used WGCNA algorithm^24^ (**Supplementary methods**; **Table S4**); however, we chose to use SpeakEasy because of its state-of-the-art performance on benchmark and real-world datasets^23^.

We validated the biological coherence of the 47 modules from 5 perspectives: (1) Gene Ontology (GO) functional enrichment analysis, (2) module coherence (“preservation”) in a separately processed set of ROSMAP samples and an entirely independent cohort, (3) concordance with co-regulation observed in epigenetic data generated from the same ROSMAP brains, (4) concordance with brain gene expression data from multiple AD mouse models, and (5) cell type-specific expression. In the first validation, we find that 29 (62%) of the modules were enriched for at least one GO functional category (Bonferroni p<0.01, **Table S5**), which is in the range of enrichments seen in other coexpression studies^14^. In the second type of validation, we assessed whether the 47 modules are preserved in: (a) a previously published DLPFC microarray dataset of 229 persons with pathologic *and* clinical AD^15^ and (b) RNA-Seq data from an independent set of 82 ROSMAP subjects. Using the Z-summary statistic that summarizes multiple measures of module preservation^25^, we observed significant evidence for preservation in 45 of the 47 modules in the independent microarray dataset and all 47 modules in the separately-processed ROSMAP subjects (Figure S5A). In a third type of validation, we assessed the robustness of these modules from a transcriptional regulatory perspective with H3K9Ac ChIP-Seq data generated from the same DLPFC samples. The H3K9Ac histone mark is found near active transcription start sites and enhancers^26^, and so coacetylation would be expected for genes that are coexpressed at the RNA level. In our H3K9Ac data, we detected for 32 of the 47 RNA-Seq-defined modules, indicating that the modules reflect, in part, the underlying epigenomic architecture (**Figure S5A**). In a fourth type of validation data, we also found that 31% of the modules are significantly preserved (Bonferroni adjustment) in gene expression data derived from the cortex of several pathology-based mouse models of AD^27^ (**Figure S5A**), which may help to prioritize those results that can be pursued further in mouse models. Finally, in a fifth type of validation we find that 13 modules are strongly enriched for cell-type specific genes, including those from microglia, astrocytes, oligodendrocytes and two neuronal subtypes (GABAergic and pyramidal neurons) (**Figure 2D**, **Table S7**) based on purified neuronal and non-neuronal cell populations, derived from the mouse brain (**Supplementary methods**). Thus, some of the modules capture previously identified cell-specific gene sets. Each of these 5 validations in parallel and orthogonal datasets indicates that gene membership of these modules is reproducible and biologically meaningful.

We next assessed the association between each module and each phenotypic trait, (**Supplementary methods**). Overall, 11 modules were associated (Bonferroni adjusted at module level, p<10^−3^) with cognitive decline, AD dementia, or AD pathology traits (**Figure 2B**). Consistent with the gene-level results, cognitive decline was associated with the greatest number of modules. In general, modules enriched for GO categories related to immunity, mitochondria, cell cycle and transcriptional regulation show a positive correlation with β-amyloid, AD and cognitive decline, while modules for neuronal or synaptic function showed a negative correlation. To replicate the phenotypic associations of these modules, we projected them onto the largest previous AD brain microarray study^28^ (the “Zhang study”) (**Supplementary methods**). Only a pathologically confirmed diagnosis of AD dementia is available in the Zhang study as an outcome, and we observed a strong concordance in both the strength and direction of the module-pathologic AD associations that we defined in ROSMAP subjects (**Figure 2C**). Notably, we observed that the module-level trait associations were more similar between these two datasets than were the univariate gene-level trait associations, further reinforcing the utility of the module-level approach (**Figure S5B**).

Because variation in cell type proportions across individuals can drive coexpression patterns between genes, some module-trait associations may be due to changes in cell type proportions that occur over the course of the disease (**Figure S3A-B**). Importantly, as we describe next, the identification of “cell type-specific modules” enabled us to guide functional validation efforts and to comprehensively model and account for changes in cell type composition that may drive large-scale transcriptomic changes in AD at the bulk tissue level.

### Identifying modules associated with disease in networks of the aging human brain

To separate a small number of direct module-trait associations from the larger number of indirect module-trait correlations, including those that may represent cell-type composition differences, we used Bayesian network (BN) inference^29^. A BN models the joint probabilities of a set of random variables as a directed acyclic graph (DAG). Here, the random variables represent module expression levels and trait values across individuals. Edges in a BN represent direct conditional dependencies between two variables: an arrow from X and Y in a BN indicates that a value taken by variable Y depends on the value taken by variable X, conditioned on all the other variables in the BN (see **Supplementary methods**).

To limit the network size for more accurate inference, only modules associated with any of the three main AD-related traits (β-amyloid load, tau tangle density, and cognitive decline) were included. We also included four modules representing the proportions of four major brain cell types - microglia, astrocytes, oligodendrocytes, and neurons - to account for the effects of changes in cell population frequencies (**Supplementary methods**). In summary, the resulting BN consisted of 18 nodes, including 11 nodes representing trait-associated modules, 3 trait nodes, and 4 “cell type modules” (**Figure 3A**).

**Fig 3.**
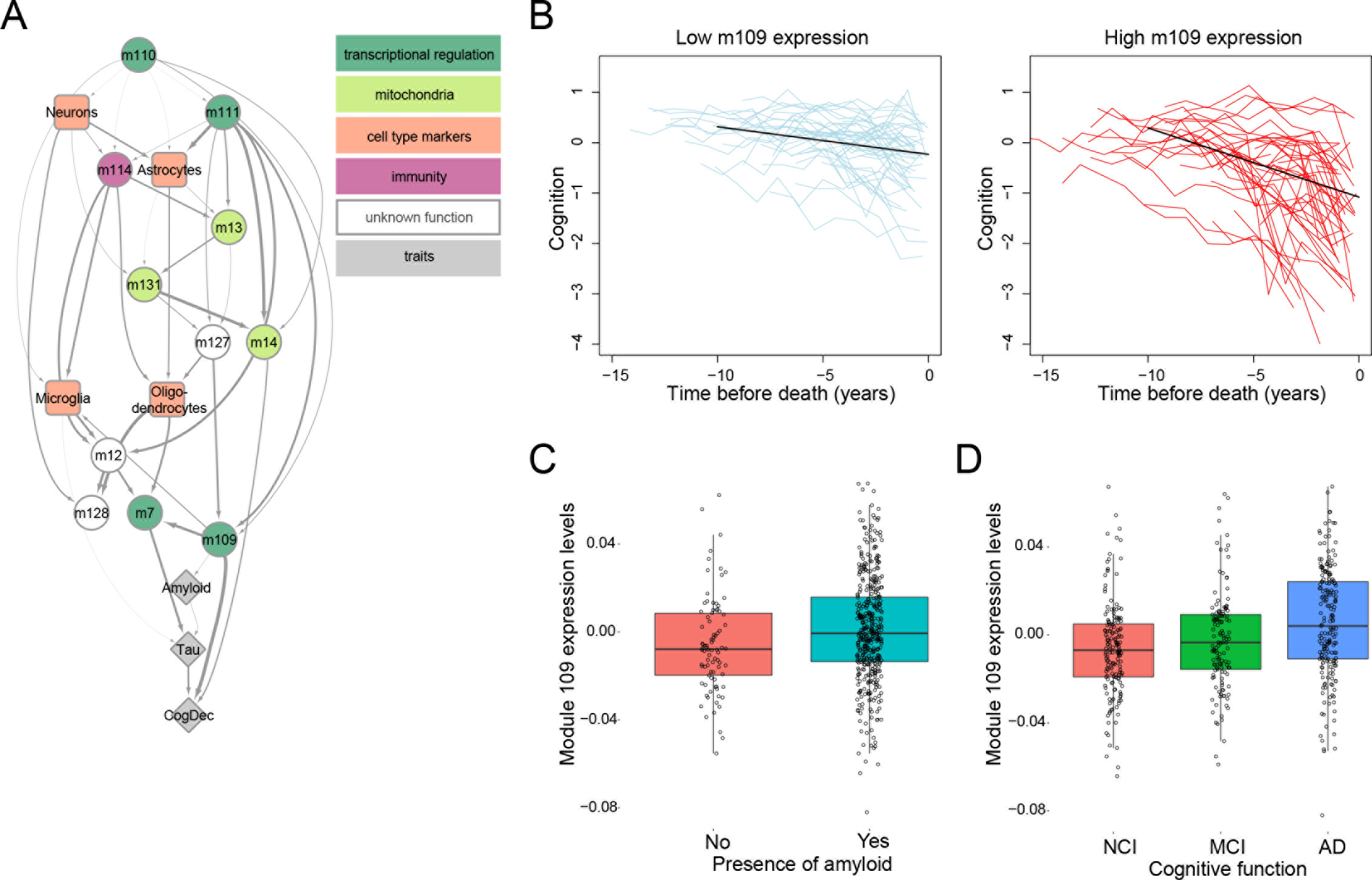

As shown in **Figure 3A**, module 109 (m109) is the module most strongly associated with cognitive decline, conditioned on all other correlated modules and modules that represent cell type proportions. It consists of 390 genes with diverse functions. Prominent functions that are enriched in this module relate to the regulation of the cell cycle and chromatin modification (**Table S5**), although there are also smaller groups of genes that are members of classical signal transduction (NOTCH) and cell-to-cell communication pathways (e.g., members of TGFβ signalling cascade). The association of m109 with cognitive decline replicates in the independently processed set of 82 ROSMAP subjects (p=0.006).

### Relation of RNA derived modules to the genetic architecture of AD

The *APOE* ɛ4 haplotype has a unique role in the disease given its large effect size (odds ratio > 3 for one copy of the *APOE* ɛ4 allele) and substantial frequency in human populations^30^. Therefore, we evaluated its effect on m109 and found that *APOE ɛ4* has a modest association with higher m109 expression (nominal p=0.03, Wilcoxon test, **Figure S6**), consistent with this haplotype’s known associations with accumulation of β-amyloid pathology and cognitive decline^31^. In a mediation analysis, after accounting for the *APOE ɛ4* haplotype, m109 remains strongly associated with both pathological AD (effect magnitude reduced by 13%, adjusted p=0.0027) and cognitive decline (effect magnitude reduced by 7.4%, adjusted p=5.03x10^−10^), implying that the effect of m109 is largely independent of this susceptibility haplotype. We extended our genetic analysis of m109 to other common AD susceptibility variants^4^ which, in earlier studies, displayed very little or no association with neuropathologic features of AD^32,33^. The 21 common AD variants did not associate with m109 (i.e., trans-eQTL analysis), consistent with the observed sparsity of trans-associations (trans-eQTLs) reported in studies with larger sample sizes^34^.

We also assessed whether any of the modules are enriched for genes found in the vicinity of AD variants (nearest genes, as previously reported^4^) and found one enriched module, m116, (p=0.0018 using the INRICH algorithm^35^) which mostly contains microglial genes (**Table S7**). However, m116 is not directly associated with cognitive decline, AD pathology, or AD dementia in this dataset (**Table S6**). But it is associated with age (p=0.003), highlighting the important role of advancing age in AD susceptibility and the fact that AD GWAS studies use a clinical diagnosis of AD and often younger control subjects to achieve very large sample sizes.

Finally, for completeness, we tested the association between the 21 common AD susceptibility alleles^4^ and the 47 modules (i.e., a module-QTL analysis) and found no significant associations after multiple testing correction, consistent with the observation noted earlier that, while these susceptibility alleles are robust risk factors, they capture only a small fraction of the variance in these AD-related traits.

### Prioritizing genes in module 109 and testing their effect on extracellular β-amyloid levels

Because of m109’s strong direct association with cognitive decline (p<10^−9^) we elected to focus our validation effort on this module. In addition to cognitive decline, m109 is associated with β-amyloid pathology (p<0.0001) (**Figure 3A and D, Figure 2B**), but we note that a mediation analysis indicated that the effect of m109 on cognitive decline is not fully mediated by the accumulation of β-amyloid. That is, m109 influences cognitive decline through β-amyloid and non-β-amyloid processes. As we describe below, due to the absence of a cellular model of cognitive decline, we focused our validation effort on m109’s association with *β*-amyloid load.

To prioritize genes in m109 for functional validation, we used several criteria: gene network connectivity, sufficient expression levels in cultured human astrocytes and/or induced human neurons (our two experimental systems), the magnitude of the gene-level association with our three AD-related phenotypes, and existing knowledge about gene function. We identified 21 genes within m109 that satisfied these criteria and were selected for experimental perturbation (see **Supplementary methods**).

For the 21 selected genes, an average of 5 shRNA constructs targeting each selected gene were tested for their knock down of gene expression (**Table S9**). To be included in our functional screen, genes had to have at least two shRNA constructs meeting our knockdown efficacy criteria of >50% reduction in expression in the target cell type. 12 genes (with 37 shRNA constructs) met this criterion in neurons and 14 genes (with 41 shRNA constructs) met this criterion in astrocytes (**Table S10**).

We performed 78 shRNA experiments to knock down (KD) the selected 14 genes in astrocytes and 12 genes in neurons. With these KD experiments, we measured extracellular levels of the pathogenic β-amyloid Aβ42 peptide, that can be readily assayed *in vitro* and is related to the defining pathologic lesion in AD (**Figure 3A**): we measured it in conditioned media from astrocyte cultures as well as iPSC-derived neuronal cultures following gene perturbation (see **Supplementary methods**). These experiments included three types of negative controls, including empty vectors and vectors that only contained GFP (see **Supplementary methods**). Since m109 is positively correlated with β-amyloid burden (**Table S2, Table S6**), the MTN approach predicts that a knockdown of expression would result in reduced β-amyloid levels.

Using an ANOVA model that accounted for testing multiple constructs per gene (**Supplementary methods**), we evaluated the effect of each shRNA construct on extracellular Aβ42 levels in contrast to the negative controls. In neurons, this outcome measure was not altered. By contrast, we identified two shRNA constructs targeting different genes, *INPPL1* and *PLXNB1*, that exceeded the Bonferroni threshold of significance (p< 0.0012) in astrocytes (**Figure 5A**; **Table S11**) in our discovery screen. Two additional constructs for these genes were found to meet a suggestive threshold (p<0.024, defined as one over the number of tests). Notably, these two genes are predicted to be upstream of the other tested genes in our BN model (**Figure 4A**), are directly connected to one another and are two of the major hubs in the m109 coexpression network (**Figure 4B**).

**Fig 4.**
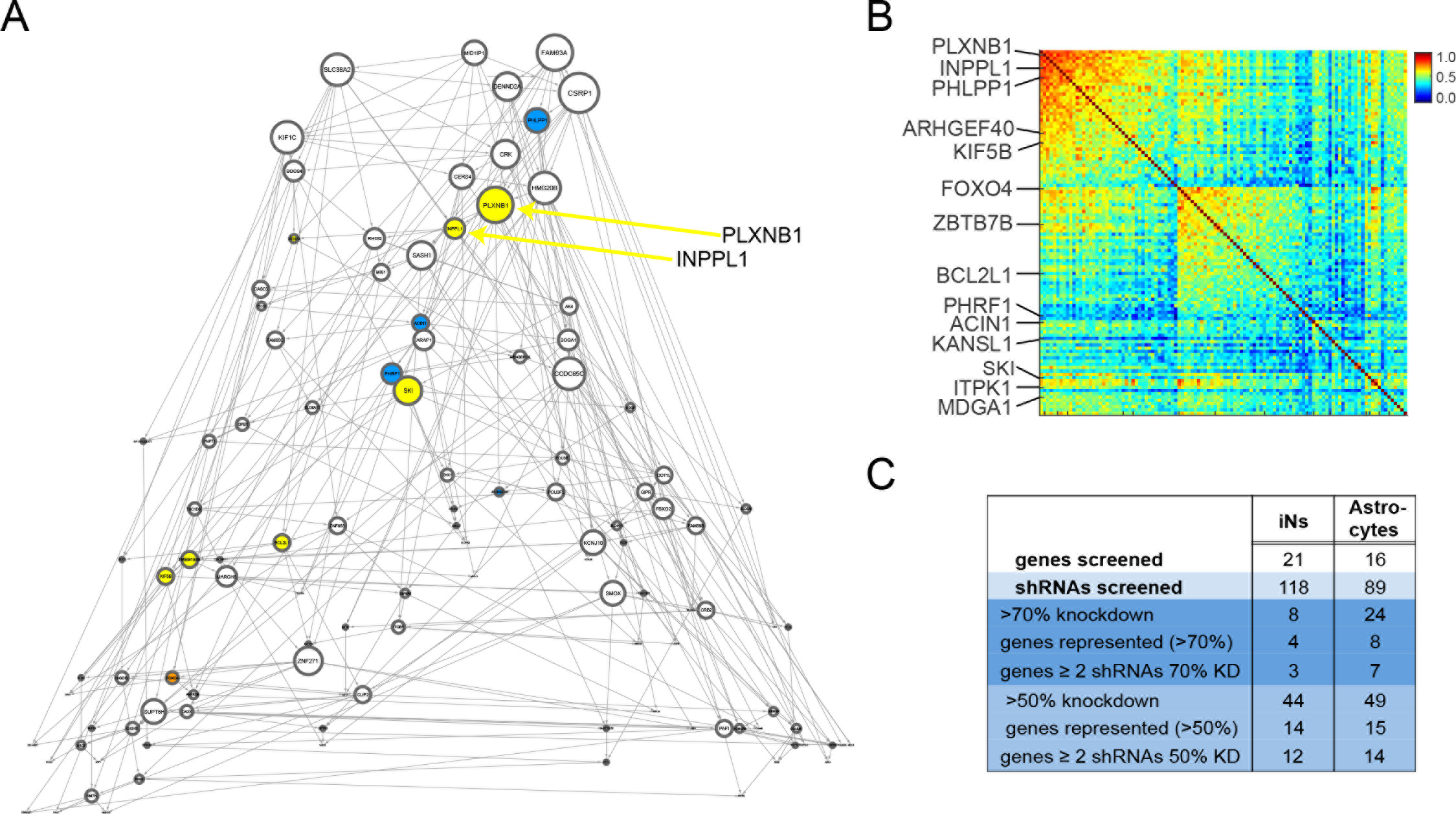

**Fig 5.**
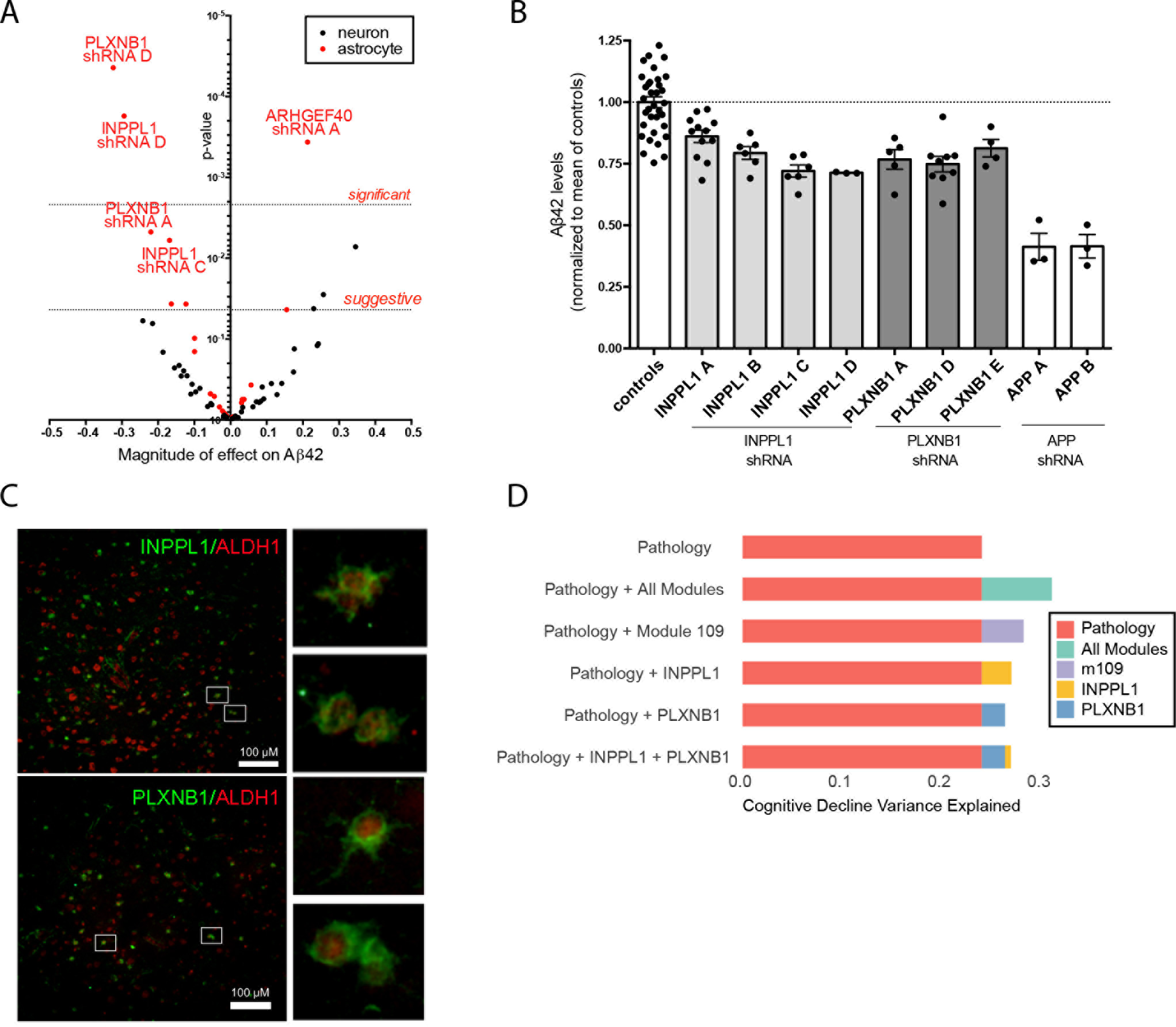

To replicate these findings, we repeated the perturbation experiments, this time including 4 shRNA constructs for *INPPL1* and 3 shRNA constructs for *PLXNB1*, as well as several negative and positive controls (see **Supplementary methods**). Specifically, as positive control we included shRNA constructs targeting *APP* the parent protein of the A*β*42 peptide). In these experiments, knockdown of both genes led to reduced extracellular levels of Aβ42 in astrocyte cultures (p_*INPPL1*_ = 2×10^−4^ and p_PLXNB1_ = 9×10^−6^) (**Figure 5B**). Overall, these results are consistent with the direction of m109 association where higher m109 expression is seen with a greater β-amyloid load. We also immunostained frontal cortex from subjects with pathologic AD and showed that both *INPPL1* and *PLXNB1* were expressed at the protein level in astrocytes (**Figure 5C**), confirming that these two genes were expressed *in vivo* in the human cell type used in the validation experiments. The astrocytes expressing these two genes are found in the vicinity of *β*-amyloid deposits (data not shown). These genes also are expressed in other cell types, such as neurons, and we cannot rule out that they may be implicated in AD in more than one cell type.

Having functionally validated a role modulating extracellular levels of pathogenic Aβ42 for these two m109 genes *in vitro*, we returned to our transcriptomic data to evaluate the magnitude of the effect of *INPPL1 and PLXNB1* on β-amyloid load. Separately, they each account for a small proportion of the variance in β-amyloid load - 2.8% for *INPPL1 and* 3.1% for *PLXNB1* - and, as is anticipated from the network model, are largely redundant as together they explain 3.1% (adjusted R^2^) of the variance in β-amyloid load (**Figure 5D**). This compares to <1% variance of this trait explained by validated AD SNPs other than *APOE.* For example, the *CR1* susceptibility allele explains 0.39% of variance in β-amyloid load in our data^36^. The effect of *INPPL1* and *PLXNB1* is somewhat stronger on cognitive decline: 5.5% of variance is explained by *INPPL1*, 4.4% for *PLXNB1*, and 5.4% (adjusted R^2^) for both. More broadly, we assessed the degree to which these two genes capture the effect of the entire module. The m109 meta-feature explains 4.3% variance in β-amyloid burden and 8.5% variance in cognitive decline, and, after accounting for *INPPL1 and PLXNB1*, we see that some of the effect of m109 remains for β-amyloid (0.95% variance, p=0.029), and more of the effect on cognitive decline persists (3.5% variance, p=2.7×10). Thus, while *INPPL1* and *PLXNB1* play an important role in m109, they do not appear to account for the effects of the entire module, suggesting that additional validation work will be needed to identify additional driver genes for m109 and that they are likely driven by non-β-amyloid processes.

## Discussion

We proposed MTN, a network-based approach, for identifying coherent biological processes and specific genes associated with multiple AD-relevant traits. A key feature of our approach is the identification of unique, direct molecular-pathological-clinical relationships, which should reduce efforts spent on spurious disease associations and on indirect associations. Further, we apply MTN to a cohort with measures of change in cognition over time, which is the most relevant clinical outcome measure of AD clinical trials for both prodromal AD as well as MCI due to AD and AD dementia. This framework and our data allowed us to identify cellular processes in the human cortex that directly relate to cognitive decline, separate from those genes that directly influence the accumulation of AD pathology. These processes are different and complementary to those identified by GWAS-derived genes, and, in the case of m109, we see how they interact with key genetic risk factors such as *APOE ɛ4*.

The central finding of this project is the existence of a robust set of coexpressed genes, supported by multiple other datasets, which is related to both β-amyloid burden and to the slope of cognitive decline in older individuals. Since modeling cognitive decline *in vitro* is challenging, we explored the relationship of some predicted influential genes in this system with β-amyloid biology. Two of these genes *INPPL1* and *PLXNB1* showed relationships to extracellular β-amyloid levels in astrocyte cultures. However, we note that the overall conclusions of this study are strongest at the systems level, and more *in vitro* experiments and careful selection of the model system will be needed to test the regulatory structure of this system as a whole.

PLXNB1 is a member of a family of proteins that mediate Semaphorin signalling, which plays a role in a number of neuronal processes including neurite outgrowth, remodelling and synaptic plasticity^37^. However, little is known about its potential contribution to cognitive decline and AD. INPPL1, also called SHIP2, is a lipid phosphatase that regulates the levels of the important second messenger PIP_3_. Levels of PIP_3_ in turn regulate downstream AKT and GSK3β signalling pathways, and *AKT* is also a member of the m109 module. *INPPL1* plays an important role in insulin signalling, and mutations and polymorphisms in the *INPPL1* gene are associated with type 2 diabetes mellitus (T2D)^38^. Transgenic mice overexpressing *INPPL1* show a disruption in insulin/IGF1 signalling mediated via *AKT*, and, interestingly, these mice show impaired memory, as assayed by three behavioral tests^39^. Further, administration of an *INPPL1* inhibitor to a mouse model of diabetes was able to rescue the synaptic plasticity and memory defects observed in this model. Thus, these *in vivo* experiments are consistent with the direction of effect that we found in our network model and in our validation experiments.

The inositol phosphatase *SHIP1* (*INPP5D*) that is structurally related to *INPPL1* harbors an AD GWAS hit^4^ and is associated with pathologic AD in ROSMAP^33^, although that gene may be more related to microglia signaling^40^. In addition, in prior work we found evidence of decreased response to insulin growth factor 1 (*IGF1*) in brains from persons with mild cognitive impairment and AD dementia relative to controls, regardless of diabetes status.^41^ Specifically, levels of postranslationally modified forms of the IGF1 receptor, IRS-1 pS^616^ and IRS-1 pS^636^/^639^, and their activated kinases were positively related to β-amyloid and negatively related to episodic and working memory, after adjusting for AD pathology. In line with these findings, our study provides evidence that *INPPL1* plays a role in cognitive decline and β-amyloid accumulation, and it opens new avenues for study of the important linkages between phosphoinositides, insulin signalling, and AD.

The MTN method has therefore identified two genes with compelling information from this and prior studies to support their further evaluation as AD target genes. It is however, important to note the limitations of our approach. First, assessing the false positive rate of MTN or other similar methods would require experiments on a large set of predicted null targets, which is currently cost prohibitive. Also, the MTN framework acts to increase network accuracy by modeling networks at two resolutions (a zoomed-out module/trait network and a zoomed-in gene network within selected modules), but increased accuracy comes at a price of including only a subset of modules in the inference process, resulting in potential loss of information. Additionally, an important “biological” limitation is functionally screening m109 genes for effects on *in vitro* β-amyloid extracellular levels, when their strongest effects were on cognitive function. This compromise thus interrogates only part of m109’s function.

In summary, we completed the initial arc of molecular network discovery and identification and validation of specific genes that are candidates for further validation and hopefully therapeutic development. This report is an initial blueprint for a quantitative systems s investigating the biology and function of the human brain.

## Acknowledgements

We thank the participants of ROS and MAP for their essential contributions and the gift of their brain to these projects. All subjects gave informed consent. This work has been supported by many different NIH grants: U01AG046152, R01AG036836, P30AG10161, R01AG015819, R01AG017917, R01AG036547. This work was done as part of the National Institute of Aging's Accelerating Medicines Partnership for AD (AMP-AD).

## Supplementary Material

1. Supplementary methods
2. Supplementary Figures S1-S6
3. Supplementary Table S1-S11 as sheets for one excel file

## Supplementary Methods and Materials

### ROSMAP pathology and clinical traits

#### Amyloid and Tau

To quantify an estimate of the burden of parenchymal deposition of beta-amyloid and the density of abnormally phosphorylated tau-positive neurofibrillary tangles levels present in the cortex at death (which we refer to as β-amyloid and tau-tangles, respectively), tissue was dissected from eight regions of the brain: the hippocampus, entorhinal cortex, anterior cingulate cortex, midfrontal cortex, superior frontal cortex, inferior temporal cortex, angular gyrus, and calcarine cortex. 20μm sections from each region was stained with antibodies to the amyloid beta protein and the tau protein, and quantified with image analysis and stereology, as previously described^1–4^. Briefly, amyloid beta was labeled with an N-terminus-directed monoclonal antibody (10D5; Elan, Dublin, Ireland; 1:1,000). Immunohistochemistry was performed using diaminobenzidine as the reporter, with 2.5% nickel sulfate to enhance immunoreaction product contrast. Between 20 and 90 video images of stained sections were sampled and processed to determine the average percent area positive for amyloid beta. PHFtau was labeled with an antibody specific for phosphorylated tau (AT8; Innogenetics, San Ramon, CA; 1:1,000). Between 120 and 700 grid interactions were sampled and processed, using the stereological mapping station, to determine the average density (per mm^2^) of PHFtau tangles. The scores across the eight regions were averaged, for amyloid and tau separately, to create a single summary measure for each protein. To create approximately normal distributions and facilitate statistical comparisons, we analyzed the square root of these two summary measures.

#### Cognitive Decline

The ROS and MAP methods of assessing cognition have been extensively summarized in previous publications^5–9^. Uniform structured clinical evaluations, including a comprehensive cognitive assessment, are administered annually to the ROS and MAP participants. Scores from 17 cognitive performance tests common in both studies were used to obtain a summary measure for global cognition as well as measures for five cognitive domains of episodic memory, visuospatial ability, perceptual speed, semantic memory, and working memory. The summary measure for global cognition is calculated by averaging the standardized scores of the 17 tests, and the summary measure for each domain is calculated similarly by averaging the standardized scores of the tests specific to that domain. To obtain a measurement of cognitive decline, the annual global cognitive scores are modeled longitudinally with a mixed effects model, adjusting for age, sex and education, providing person specific random slopes of decline. The random slope of each subject captures the individual rate of cognitive decline after adjusting for age, sex, and education. Further details of the statistical methodology have been previously described^10^.

#### Pathologic diagnosis of AD

A pathologic diagnosis of AD was determined by a board certified neuropathologist blinded to age and all clinical data and using modified Bielschowsky silver stained 6 micron sections of hippocampus, entorhinal cortex, midfrontal cortex, midtemporal cortex and inferior parietal cortex. The diagnosis follows the recommendation of the National Institute on Aging-Reagan criteria^11^. Briefly, based on the scores of Braak stage for severity of neurofibrillary tangles and CERAD estimate for burden of neuritic plaques, a pathologic AD diagnosis requires an intermediate likelihood AD (i.e., at least Braak stage 3 or 4 and CERAD moderate plaques) or a high likelihood AD (i.e., at least Braak stage 5 or 6 and CERAD frequent plaques).

#### Clinical diagnosis of AD at the time of death

Annual clinical diagnosis of AD dementia follows the recommendation of the joint working group of the National Institute of Neurological and Communicative Disorders and Stroke and the AD and Related Disorders Association^12^. The diagnosis requires a history of cognitive decline and evidence of impairment in memory and at least one other cognitive domain. After a participant had died, a neurologist specializing in dementia reviews all available clinical information and provides a summary opinion with regards to the most likely clinical diagnosis at the time of death. The summary diagnosis was blinded to all neuropathologic data, and case conference are held for consensus as necessary^13^. AD dementia includes persons with probable or possible AD dementia, i.e. AD dementia with or without comorbid conditions that may be affecting cognition.

### ROSMAP RNA-Seq data quantification and QC

RNA was sequenced from the gray matter of dorsal lateral prefrontal cortex (DLPFC) of 542 samples, corresponding to 540 unique brains. These samples were extracted using Qiagen's miRNeasey mini kit (cat. no. 217004) and the RNase free DNase Set (cat. no. 79254). RNA was quantified using Nanodrop. Quality of RNA was evaluated by the Agilent Bioanalyzer. All samples were chosen to pass two initial quality filters: RNA integrity (RIN) score >5 and quantity threshold of 5 ug (and were selected from a larger set of 724 samples). RNA-Seq library preparation was performed using the strand specific dUTP method^14^ with poly-A selection^15^. Sequencing was performed on the Illumina HiSeq with 101 bp paired-end reads and achieved coverage of 150M reads of the first 12 samples. These 12 samples will serve as a deep coverage reference and included 2 males and 2 females of non-impaired, mild cognitive impaired, and Alzheimer's cases. The remaining samples were sequenced with a target coverage of 50M reads; the mean coverage for the samples passing QC is 95 million reads (median 90 million reads). The libraries were constructed and pooled according to the RIN scores such that similar RIN scores would be pooled together. Varying RIN scores results in a larger spread of insert sizes during library construction and leads to uneven coverage distribution throughout the pool.

RNA-Seq data were processed by our parallelized pipeline. This pipeline includes trimming the beginning and ending bases from each read, identifying and trimming adapter sequences from reads, detecting and removing rRNA reads, and aligning reads to reference genome. The non-gapped aligner Bowtie was used to align reads to the transcriptome reference^16^, and RSEM was used to estimate expression levels for all transcripts^17^. The FPKM values were the outcome of our data RNA-Seq pipeline.

To remove outlier samples based on quantified expression profiles, following a previous approach^18^, the D-statistic was computed as the median correlation of all genes (based on expression profiles) of each sample with all other samples. An additional 13 samples with D-statistics < 0.9 were deemed outliers and excluded.

For the 2 individuals with replicate samples, we only took (at random) one of the replicates, and excluded 19 samples without genotyping data. Thus 508 unique samples were analyzed for derivation of gene modules. For trait analysis, we further excluded 30 samples where clinical and/or pathological assessment was not complete, resulting in 478 brain samples (unique individuals) analyzed with respect to AD-traits.

### RNA-Seq data normalization

We applied quantile normalization to FPKM first and then used the combat algorithm^19^ to remove potential batch effect. Expression levels were quantified for 55,889 unique genes and 190,051 transcripts. We placed a threshold for expression, only keeping gene’s with at least 4 reads in 100 individuals (yielding 13,484 significantly expressed genes).

After quantile normalization and batch correction, we used linear regression (on log2 expression data) to remove the effect of major biological and technical confounding factors on a per-gene basis. Biological confounding factors include three genotyping PCs (to represent ancestry), age at death, and sex. Technical confounding factors include RIN, number of risbosomal bases, number of aligned reads, study index (ROS or MAP), and Post Mortem Interval (PMI).

In addition to known confounding factors, previous studies have shown that in some cases removal of “hidden” confounding factors (e.g., “surrogate variables”^20^) could increase statistical power (e.g., the consistent reported increase in statistical power for cis eQTL studies^21^. These hidden factors can be estimated by the top expression PCs (or factors)^22–24^. Here, for reasons described below, we did not find sufficient evidence to normalize for potential hidden confounds.

First, we hypothesized that if the removal of expression PCs does indeed improves the identification of disease associated genes (either through reducing false positives, or increasing statistical power), then as we remove expression PCs, we should see an increase in the strength of the association between known AD genes and AD traits. Therefore, we assessed the differential expression of 32 known AD genes (from literature and GWAS) after removing 1 up to 10 expression PCs. As shown in **Figure S1A**, we noted that removal of PCs generally degraded the association strengths. Second, we noted that the top expression PCs correlate with known cell type markers. Therefore, without explicitly adjusting for expression PCs, some of the estimated modules represent patterns that are predictive of cell type proportions (**Tables S6**). As described in the main manuscript, our approach uses these “cell-specific” modules to correct for cell type composition changes - we reasoned that for a cell-specific module, the relative expression across individuals is indicative of proportion of the represented cell type. We think that this post-hoc approach for assessing cell type proportion presents a powerful method for identifying interesting cell types and conservatively correct for cell type proportion changes that may lead to differential expression. Third, we noted that after removing the top few PCs, clustering algorithms tended to produce many small clusters/modules (modules with few gene members). Thus, we chose to only account for “known” covariates and not any hidden covariates. Hidden covariates are captured by broad modules that represent cell types, and they are analyzed in a post-hoc manner.

### Association analysis

#### TWAS analysis

For continuous traits (β-amyloid, tau-tangles, and cognitive decline), we used spearman rank correlation (and the associated p-values) to quantify the strength of associations between each gene’s (or transcript) expression levels and a given trait (**Table S2**). We used a t-test to assess the association between binary traits (diagnosis status) and gene expression levels. We noted that tau pathology was associated with a much smaller set of genes than β-amyloid pathology.

#### Module-to-trait association analysis

Each module (see below for module derivation) is first represented by the mean expression level of all genes assigned to that module (expression data is standardized prior to computing module means). Then, the module’s mean expression vector is associated with each AD-trait, using the Spearman correlation for continuous traits and Pearson correlation (whose pvalue is equivalent to a t-test) for binary traits.

### Deriving gene modules and assessing their quality

We used the SpeakEasy (SE) consensus clustering algorithm^25^ to derive gene modules from normalized gene expression data. We quantify the consensus clustering results from 100 initializations of the SE algorithm, which produced 257 modules, 47 of which contained at least 20 gene members (meaning modules with >20 members are assigned for 98% of genes) and were examined in downstream analyses (visually summarized in **Figure S4**). The assignments of gene to modules are provided in Supplementary **Table S3**.

We used the SE algorithm as it was recently shown to be the state-of-the-art in terms of identifying robust clusters in the Lancichinetti-Fortunato-Radicchi (LFR) benchmarks, as well as in a wide range of biological networks. Specifically, on these previous benchmark experiments, the SE algorithm outperformed standard hierarchical clustering algorithms and their derivatives such as WGCNA^26^. But on the ROSMAP RNA-seq data, we also investigated the degree to which the clustering solutions of SE overlaps with those produced by the WGCNA^26^ algorithm. Because individual cluster-to-cluster comparison metrics can be highly unstable, we compare the overall similarity across all clusters identified by SE and WGCNA collectively (**Table S4**). To make the comparison even more robust, we utilized multiple proposed measures of partition similarity^27^. These results indicate a significant similarity between SE and WGCNA, but to a lesser extent than variation between WGCNA runs under slightly different parameter setting. Thus, our clustering results carry a similarity to those of the previous WGCNA method, but based on performance benchmark comparisons and a wide range of real-world networks, are likely more accurate.

#### Module functional enrichment analysis

For each module and gene set, we computed a hypergeometric enrichment p-value for enrichment in any gene ontology category, and we used a Bonferroni-adjusted p-value of <0.01 to identify significantly enriched functions for each module. We used all DLPFC expressed genes (identified in this study) for the “background” set. According to this analysis, 29 of the 47 modules (or 62% of the modules) were significantly enriched for at least one Gene Ontology category, which covers 8368 genes (also 62% of expressed genes) (see **Table S5**).

#### Cell type composition and relationship with gene modules

Random variation in the proportion of cell types across samples (individuals) can lead to identification of modules that are enriched in genes that are highly expressed in particular cell types. To assess which of the modules have some relationship to cell types, we calculate the median rank of all genes in a module in the rank-ordered expression of genes in a given cell type (see below for sources of cell-type specific gene expression). We repeat this calculation for all combinations of modules and cell types. This provides an indicator of when modules contain genes that are highly expressed in a canonical cell type. Because modules generally consist of hundreds of genes, this test is well powered to detect even minor enrichment, with nearly every module showing significant enrichment for some cell type, down to the minimum p-value (based on 10,000 permutations). Therefore, we also provide a practical measure of enrichment effect size, which is the increase in the median rank of genes in a module in the sorted expression of genes from a given cell type, compared to the median rank from permuted gene sets. In our analysis, we use the effect size of 1.4 to identify cell-type associated modules. We chose the 1.4 threshold because we found that at this threshold, each of the major brain cell types (astrocyte, microglia, oligodendrocytes, and neurons) will be represented by at least one module. **Table S7** provides the results of this analysis.

The origin of the cell type signatures is gene expression from mouse brain, wherein cell types are better annotated with marker genes than in humans and where a larger amount of transcriptome cell type profiles are available. We downloaded 18 large appropriate datasets from GEO on the Affymetrix 430 GeneChip (microarray) platform, which were applied to measure gene expression in a range of common cell types in the mouse brain^28–45^. These cell types were captured by a range of methods, including fluorescence assisted cell sorting and laser capture microdissection. These data were often generated from disease models or exposed to various perturbations. Single-cell RNAseq is an excellent mechanism to identify cell types in an unbiased manner, but for our purpose the largest brain cell type studies in GEO do not sequence deeply enough to be useful: they show distinct expression for a few thousand genes, compared to the twenty thousand surveyed on the microarrays. To generate a genomic signature for each cell type, we simply take the median value across all replicates of a gene cell type.

For our intended use of validating the coherence of expression among our inferred modules, cell type enrichment is a useful tool that shows substantial coherence of cell type-specific expression and module membership. For purposes beyond validating the plausibility of clustering results, additional caution is needed for interpreting these results. Negative findings for a module with regard to cell type should not be seen as conclusive, due to the cross-species nature of the comparison, but positive findings with large effect sizes are likely to be meaningful. On the other hand, positive findings do not necessarily indicate the literal presence of a given cell type. Indeed some cell types, such as Bergmann cells, would not be expected to be found in the frontal cortex; however, they may utilize some subcellular system which is represented by a coexpression module derived from cortex. For instance, due to the high metabolic demands of fast-spiking PV+ (parvalbumin positive) cells have an abundance of mitochondria. Thus a module related to mitochondria may show up as enriched in genes highly expressed in PV+ cells, although that module may well be driven by mitochondria levels in a different cell type, or by mitochondrial function across cell types. In other cases, module enrichment in cell-type specific genes may literally indicate the large presence of that cell type in the frontal cortex, such as the case of module 116, which contains many genes, which when perturbed indeed result in effects in microglia^46^.

#### Replication of gene modules

We used the module preservation statistic^47^ to assess whether a given module is preserved in a second (validation) dataset (**Figure S5A**). The module preservation statistic (called “z-summary”) combines multiple metrics that assess the conservation of co-expression patterns within a given module in a second (validation) dataset. As validation data, we employed four datasets: a) microarray gene expression data from cortex (DLPFC) from a previous study by Zhang and colleagues^46^, b) microarray gene expression data from mouse brain from a previous study by Matarin and colleagues^48^, c) an independent test dataset from the ROSMAP study (unpublished, data available from Synapse doi:10.7303/syn3388564) d) histone modification data (H3K9Ac chromatin immunoprecipitation followed by sequencing, unpublished, available doi:10.7303/syn4896408). We summarize the analyses of these data below.

#### a) Microarray DLPFC dataset

Processed microarray gene expression data and meta-data were downloaded from GEO (accession GSE44772)^46^. Meta-data included a pathology variable (binary), age, gender, and brain region. Using linear regression (on log2 expression data), we removed the effect of age and gender, and analyzed 229 samples from the DLPFC region. 10,330 measured genes overlapped with genes defined as expressed in the ROSMAP dataset (this study).

#### b) Mouse brain gene expression dataset

We used the mouse gene expression datasets generated by Matarin and colleagues^48^. We only used gene expression data generated from cortex, which resulted in 112 samples, and included 38 control mice and 74 transgene mice (mutant human APP, PSEN1, APP/PSEN1, mutant human MAPT gene). For the purpose of module preservation statistic, all 112 samples were used to compute module preservations.

#### c) Module coherence in ROSMAP test samples

We used 82 samples from the ROSMAP project that were not included in the primary analysis. These samples were extracted and quantified after the initial data freeze for the primary analysis. These 82 samples were processed the same way as the samples used in the primary analysis: we accounted for the same set of technical and biological confounds in these samples using linear regression. The RNA-Seq data from these 82 samples are available from the Synapse database (reference number syn5615307).

#### d) Histone modification (acetylation) data

Histone acetylation data using ChIP-Seq (H3K9Ac chromatin immunoprecipitation followed by sequencing) is available for 712 samples. To quantify histone acetylation, after sequencing, single-end reads were aligned by the BWA algorithm^47^ against the human reference genome GRCh37. Peaks were detected in each sample separately by MACS2^48^ using the broad peak option and a q-value cutoff of 0.001. Pooled genomic DNA of seven samples was used as negative control. A combination of five ChIP-seq quality measures^49^ were employed to detect low quality samples: samples that did not reach (i) ≥ 15.10^6^ unique reads, (ii) non-redundant fraction ≥ 0.3, (iii) cross correlation ≥ 0.03, (iv) fraction of reads in peaks ≥ 0.05 and (v) ≥ 6000 peaks were removed. In total, 669 out of 712 samples passed quality control. 441 of the 669 samples overlap with RNA-seq samples. H3K9ac domains were defined by calculating all genomic regions that were detected as a peak in at least 100 (15%) of the 669 samples. Regions neighboured within 100bp were merged and very small regions of less than 100bp were removed. To map peaks to genes, we identified the closes peak to the TSS of each gene. Finally, quantified histone acetylation data was quantile normalized to account for variability in sequencing depth across individuals. To quantify the preservation of each module, we considered the “co-acetylation” signal at the closest peak to each gene. This histone modification data is publicly available form the Synapse database (reference number syn4896408).

### Replication of module-to-trait relationships

We used a previous microarray data by Zhang and colleagues (consisting of 229 samples—see above)^46^, to replicate the associations between modules and AD-traits. In particular, we used the module memberships for the 47 modules identified in the ROSMAP samples to compute the mean of each module in the Zhang et al. dataset (as above, module means are computed after standardization of gene expression data). Then, the mean expression vector for each module was associated with relevant AD-traits. As shown in **Figure 2C**, we observed that module-to-trait relationships were highly reproducible.

Using the Zhang et al., study, we also compared the replication module-level trait associations with gene-level trait associations. In particular, as a test-statistic for module-level replication, we computed the correlation between two vectors: a vector of length 47 (number of modules) representing the association between each module and the binary pathology variable in this study, and a vector of length 47 representing the association between each module and the binary pathology variable using data from the Zhang et al. study. The Spearman correlation between these two vectors is 0.9. To compare this to gene-level analysis, we performed a resampling-based analysis: we randomly sampled 47 genes (among the 10,330 genes that overlap between this study and the Zhang study) 10,000 times and computed the correlation between gene-to-trait vectors in this study and Zhang et al. study. As shown in **Figure S5B**, we observed module-level analysis was significantly more concordant between the two studies, as compared to the gene-level analysis.

### Bayesian network structure learning to compute the module-trait network

A Bayesian network (BN) is a probabilistic graphical model that represents a set of random variables and their conditional dependencies using a directed acyclic graph (DAG). Bayesian structure learning algorithms (such as the MCMC algorithm used here^49^) enable the estimation of the DAG that describes the conditional dependencies between a set of random variables in a data-driven manner. Here, we used Bayesian network structure learning to estimate the DAG that describes the relationship between three types of variables: modules, cell type proportions and AD traits. In particular, we included 11 modules in the DAG that were significantly associated with at least one of the three main AD-related traits (β-amyloid, tau-tangles, and cognitive decline), after accounting for multiple testing with the Bonferroni procedure. As described above, some of the modules are enriched for known cell-type-specific genes and thus their expression levels may present a proxy for cell type proportions (and it’s variability across individuals in this study). Therefore, we included four cell-specific modules that enable us to model cell-type specific changes. We chose these modules as the ones with the most “fold enrichment” for the four major brain cell types: neurons (m187), astrocytes (m107), microglia cells (m116), and oligodendrocytes (m123). There were several modules associated with one of the neuronal subtypes (**Figure 3D**), however, many of these modules had correlated expression profiles. We reasoned that in the absence of experimental data measuring proportion of each of the neuronal subtypes in our samples for validation, we don’t have the resolution to accurately model each of the neuronal subtypes in the BN analysis. Therefore, we chose to select the most enriched module for any of the neuronal cell types (m187) to broadly represent “proportion of neuronal cells” in the BN analysis.

The ensemble MCMC approach used here was recently shown to be among the state-of-the-art algorithms for Bayesian network structure learning when applied to gene expression data^49^. The essential difference between the REV move method we employ^50^ and classic structure learning is that for each revision of the proposed gene-gene network, the REV algorithm proposes larger changes to the evolving network structure. The benefits of these extensive updates and the specific type of network alterations is to avoid becoming trapped in a locally optimum solution - which has been demonstrated using simulated gene expression data analogous to ROSMAP^49^. Using this approach, we ran structure learning 100 times with stochastic initialization. The final Bayesian network was selected using Bayesian model averaging^51^, and we also visualize the raw edge count among the 100 models as edge width (**Figure 3A**).

### Identifying target genes in module 109

The Bayesian network structure learning framework described above identified module 109 (m109) as the module that is most strongly associated with cognitive decline and amyloid. Additionally, we confirmed that the association between m109 and amyloid is not simply explained away by variability in cell type proportions as captured by expression levels of standard cell type markers. Thus, we reasoned that m109 is the module most directly associated with cognitive decline and amyloid load in this sample. Therefore, we devised an approach to prioritize genes in m109 (which contains 390 genes) for experimental validation as described below.

First, we reduce the number of plausible genes by taking the top 112 genes (among the 390 genes assigned to m109) that are (a) most strongly associated with cognitive decline and amyloid load, and (b) are connected to other genes in an initial Bayesian network of 390 genes. Next, we construct a Bayesian network for representing conditional dependencies between these 112 genes in a DAG (**Figure 4A**). The DAG enables us to rank genes based on their number of conditional dependent connections to other genes in the module. As rationalized in previous studies^46^, we selected genes with the highest number of connections (**Table S8**) as nodes likely to influence the expression of many other genes and consequently the biological function of the module. We then further filtered these to X tested genes based on sufficient expression (FPKM>1) in the astrocyte or iNs *in vitro* model (**Table S8**) as well as the availability of efficient shRNAi constructs for knockdown experiments (**Table S9; Table S10**).

### Experimental validation of target genes

#### Cell Culture

Induced pluripotent stem cells (iPSCs) were maintained in iPSC media consisting of 400mL DMEM/F12, 100mL Knockout Serum Replacement, 5mL penicillin/streptomycin/glutamine, 5mL MEM-NEAA, and 500μL 2-mercaptoethanol (all from Invitrogen) with fresh addition of bFGF (Millipore) at 10μg/mL. Differentiation was carried out as previously reported with minor modifications (Zhang et al. 2013). iPSC-derived neurons are plated on DIV4 on matrigel coated 96-well plates (25,000 cells/well) and maintained in media consisting of 485mL Neurobasal Medium (Gibco), 5mL Glutamax, 7.5mL 20% Dextrose, 2.5mL MEM NEAA with 1:50 B27, BDNF, CNTF, GDNF, and doxycycline added just prior to use.

Human cortical astrocytes were obtained from Sciencell and cultured in basal media containing 2% FBS, 1% penicillin/streptomycin, and 1% astrocyte growth supplement (Sciencell). Astrocytes were trypsinized and plated at a density of 40,000cells/well of a 96-well plate and maintained for no more than 7 days.

#### Viral Transduction

To perturb gene targets, astrocytes were plated one day prior to transduction by 10uL of lentivirus (shRNA titer range 1.4×107-3.43×108 virus particles/mL; ORF titer range 6.82×106-4.3×107 virus particles/mL) in 40uL of growth media. iPSC-derived neurons were transduced on DIV17 with 25uL virus in 25uL growth media. After ∼18 hours, virus containing media was removed and replaced with fresh media and cells incubated for an additional ∼96 hours. Condition media was collected and stored at −20, while cells were lysed for either RNA or protein purification.

#### qPCR

Samples were prepared for qPCR using the Power SYBR Green Cells-to-Ct kit (Ambion) according to manufacturers instructions. qPCR was performed using Fast SYBR Green Master Mix (Applied Biosystems) and three technical repeats assessed with a ViiA 7 System (Applied Biosystems). Relative expression was calculated using the ΔΔCT method as described (Livak, K.J. and Schmittgen, T.D. (2001)) and GAPDH normalization.

#### Immunocytochemistry/Microscopy

Cells were fixed in 4% paraformaldehyde (Sigma) and permeabilized in 0.1% Triton X-100 in donkey serum (Jackson ImmunoResearch) prior to overnight incubation with primary antibodies. Following secondary antibody incubation for 1 hour, cells were treated with DAPI. Images of immunocytochemistry stained and fluorescence expressing cells were collected using a Zeiss LSM710 confocal microscope and ZEN black software.

#### Aβ ELISA

To determine the secreted Aβ peptide concentrations, condition media generated from cells having undergone viral transduction was subject to 6E10 Aβ peptide panel multiplex ELISA following manufacturers instructions (Meso Scale Discovery). Wells were blocked prior to 2-hour incubation of detection antibody solution with sample condition media or standard. Plates were read using an MSD SECTOR Imager 2400 and resulting peptide concentrations were normalized to total protein for each sample.

#### Statistical Analysis

Aβ42 secretion levels were measured in 6 batches for the discovery dataset, with each batch containing up to 11 control samples (see **Table S11**). These control samples consist of three different types of negative control experiments: a) no infection (“Non”), b) empty vectors (“empty”), and c) same vector as for all shRNA constructs but only includes green fluorescent protein (GFP). As these controls were not significantly different from each other, they were pooled into a single “control” group in our statistical analysis. We corrected for batch effects in two ways. First, each value was converted to “fold-change” by dividing the value measured in the cells perturbed with an shRNA by the mean of all of the controls in the corresponding batch. Second, we included indicator variables for each batch in our statistical models. The replication dataset was created in three batches, and we converted Aβ42 secretion levels into “fold-change” from the mean of the control samples as for discovery study. Fold-change was used as the independent variable in a general linear model (glm) that included all shRNAs, and we tested whether the average fold-change for each shRNA is different from 0.

**Figure S1.**
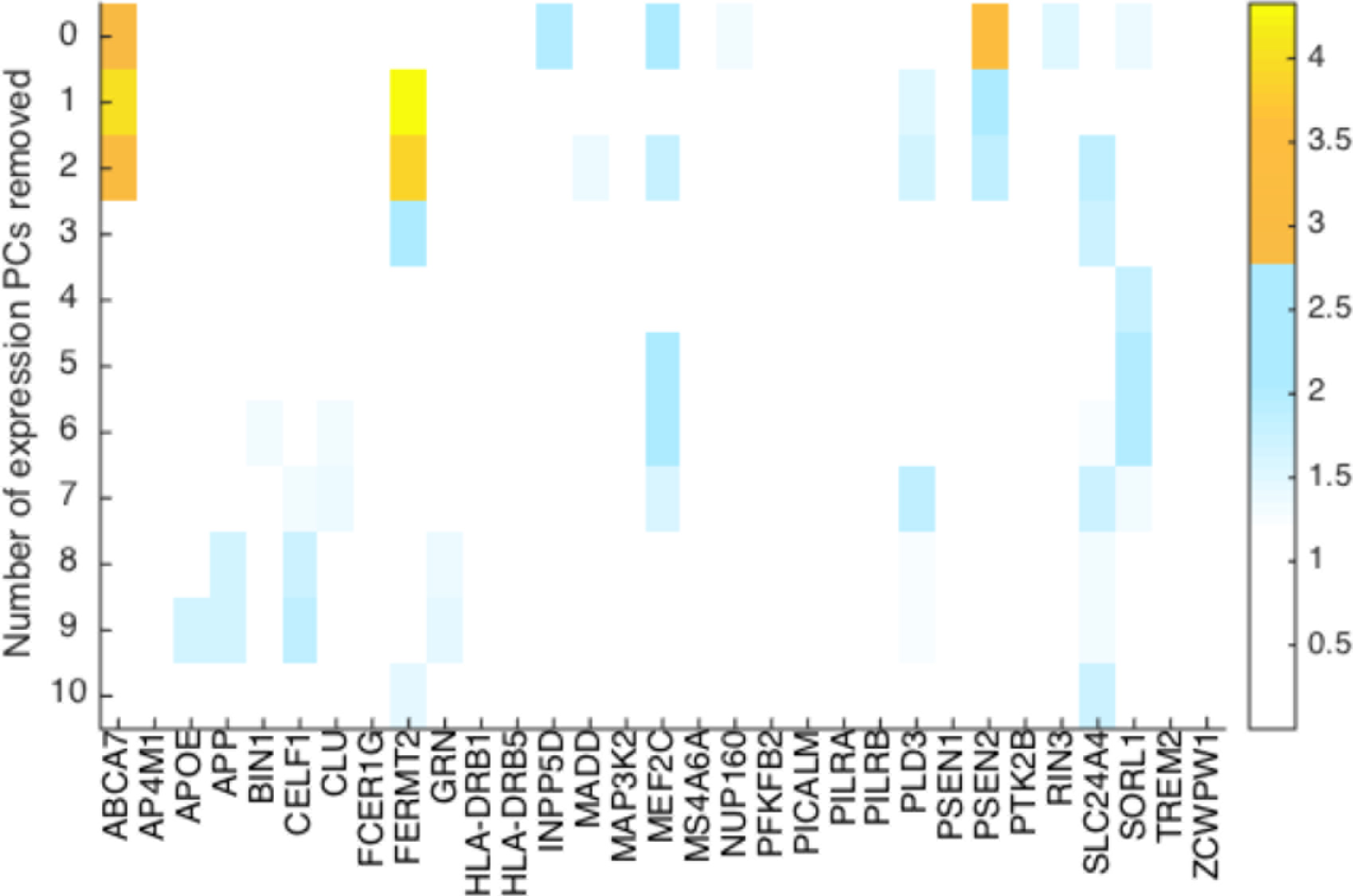
Figure shows the association strength (negative log10 pvalue) between 31 previously identified AD genes (in columns)(21 from GWAS and the remainder from smaller scale functional studies) and clinical AD. Rows report the results of the association analyses to clinical AD after removing the stated number of principal components (PCs) derived from the RNA-Seq data. The color scale reporting the level of significance is displayed to the right of the figure. This figure demonstrates that removing expression PCs typically does not improve the association strength between gene expression level and AD status.

**Figure S2.**
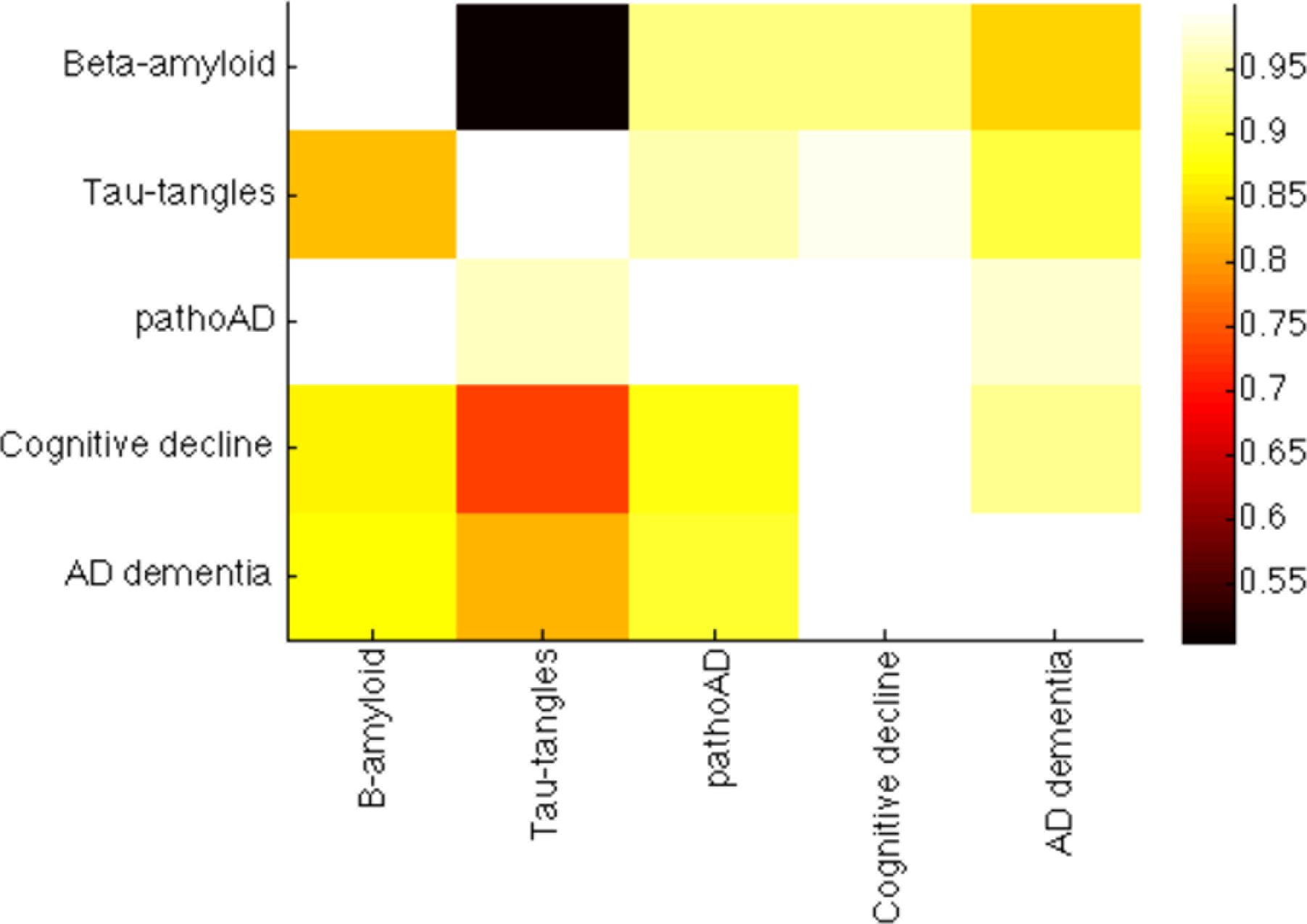
For each of the five AD-related traits shown in the figure, a univariate transcriptome-wide association study (TWAS) was conducted. To quantify the proportion of genes whose expression is associated with each pair of AD traits, the following procedure was used: for each trait i, we used the 0.05 FDR threshold to identify genes whose expression associate with trait i. In this case, trait i is the “discovery” sample. Then, we used the pi1 statistic to quantify the proportion of the 0.05 FDR genes that are deemed as true positives for trait j, and so trait j is the “replication” sample. In summary, element (i,j) in this figure represents the pi1 statistic when assessing the 0.05 FDR genes (identified in the discovery sample) from trait i (rows) in the replication sample (trait j)(columns).

**Figure S3.**
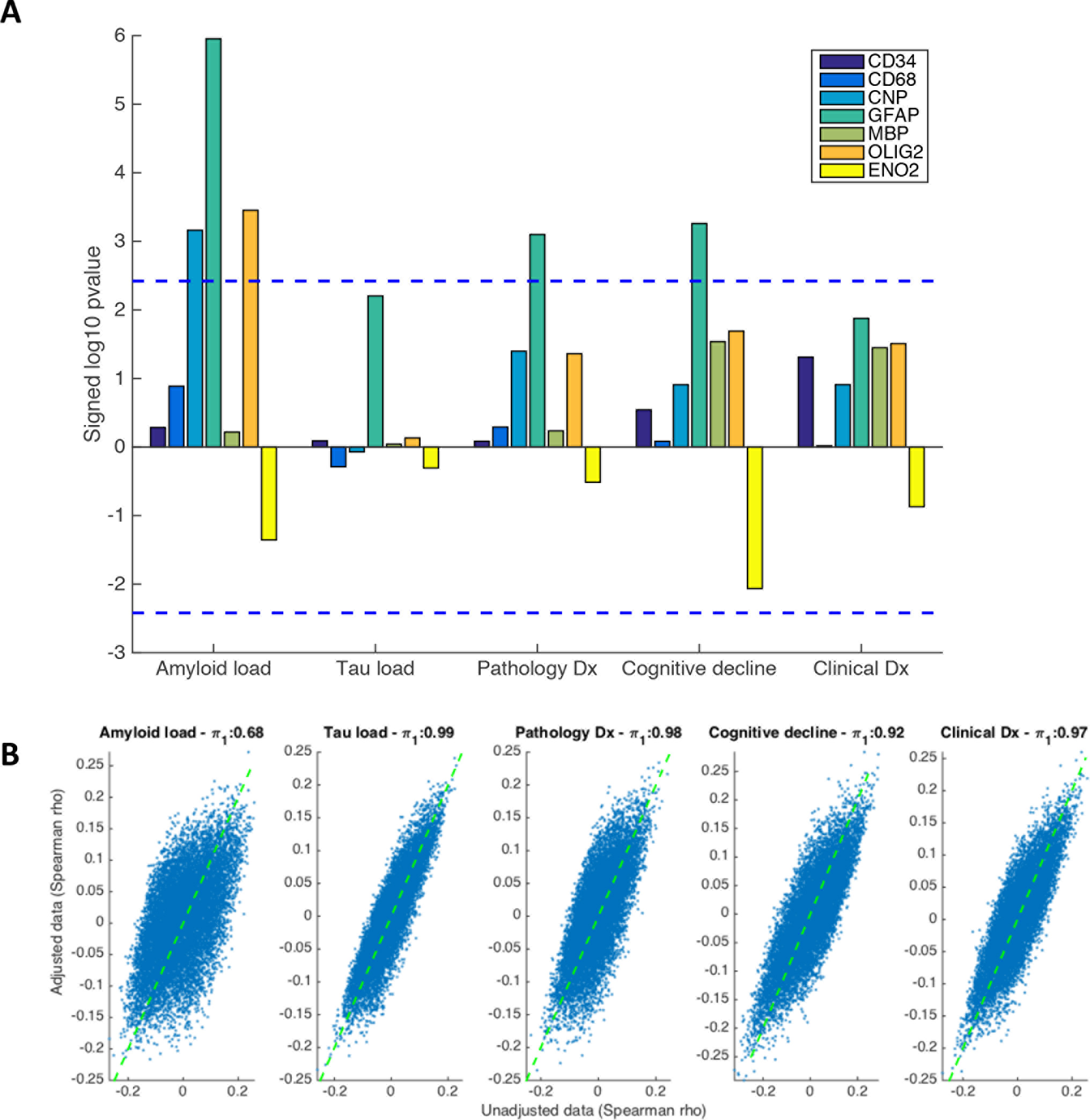
**(A)** This figure shows the correlation strength (signed, negative log10 p-value: positive denotes a direct correlation) between 7 known gene markers of common brain cell types and the AD-related traits analyzed in this study. As shown, GFAP, which is a marker of astrocytes, is most strongly associated with amyloid load. **(B)** This figure shows the correlation strength between expression level of all genes with an AD-related trait before (x-axis) and after (y-axis) adjusting for 7 standard cell type markers (shown in Figure S3A).

**Figure S4.**
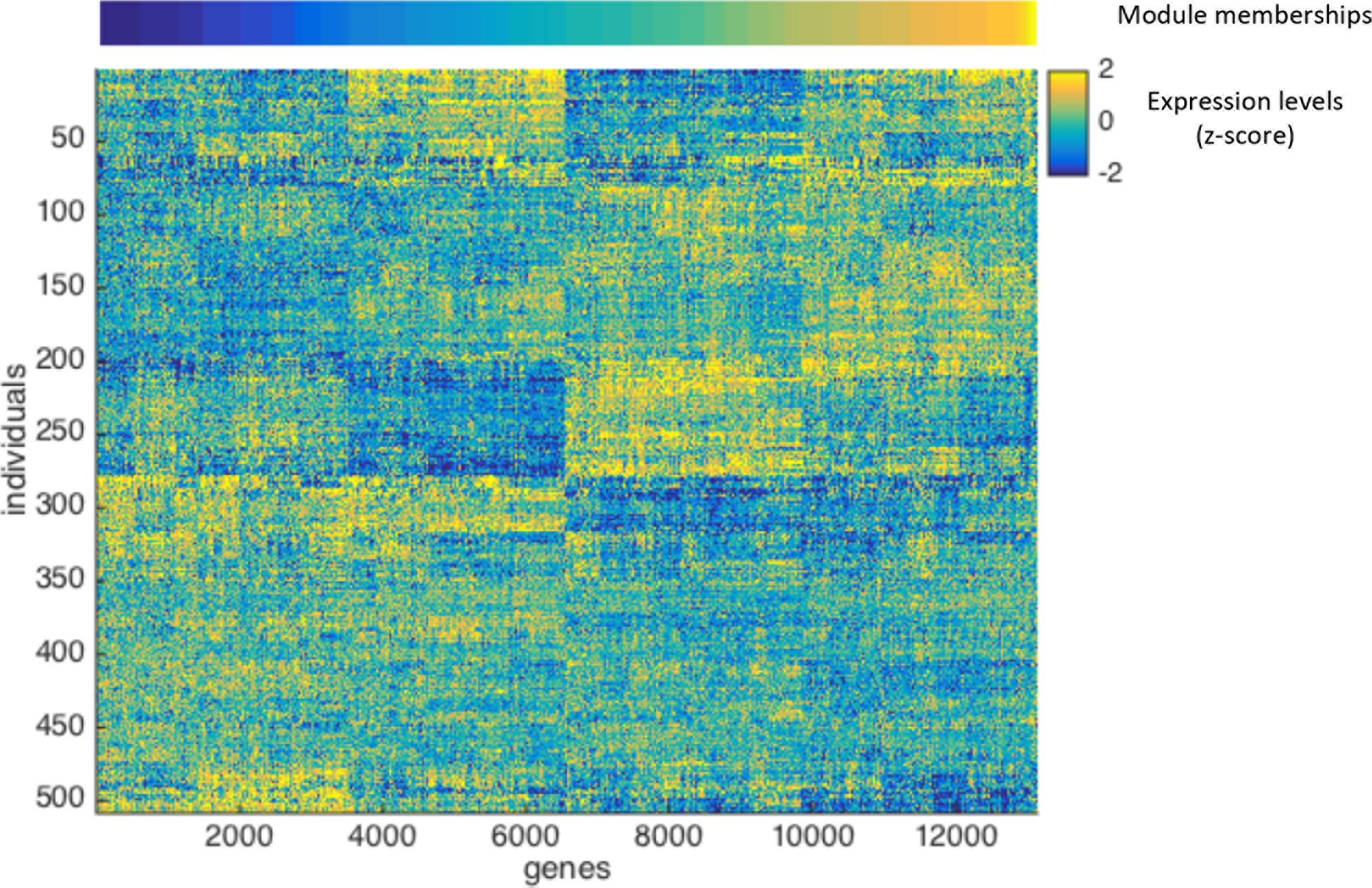
This heatmap illustrates the expression patterns of all measured genes (columns), grouped by module, in all analyzed DLPFC samples (rows). Module membership is shown at the top of the image. The color scale for relative expression is shown to be the right of the figure.

**Figure S5.**
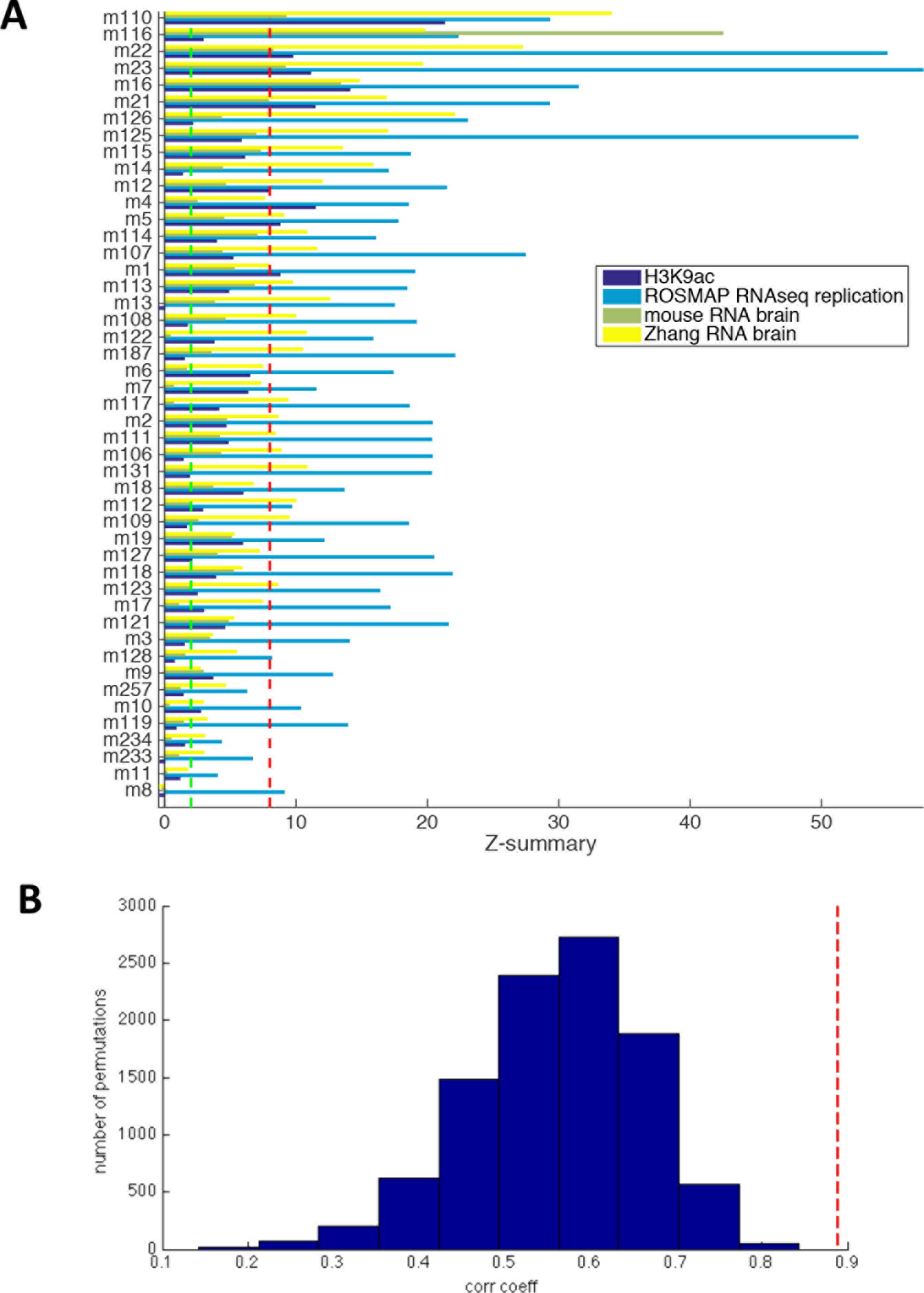
Module preservation and replication. **(A)** Module preservation (z-summary, x axis) as assessed in the 4 test datasets. Each row reports the preservation of a module in the 4 datasets. The red dashed line marks the “strongly preserved” threshold and the green dotted line marks the “moderately preserved” threshold (as defined empirically by *Langfelder, PLOS Computational Biology 2011*). **(B)** We assessed the replication of the trait to transcriptional measure associations, at the gene-level and module-level (see Supplementary material). To do so, we computed the correlation between vectors of module-to-trait association in our study and in the Zhang et al. study (shown by the red dotted line). We compared the observed correlation between module-trait vectors with gene level associations through resampling. The histogram shows the empirical distribution of correlation coefficient between vectors of gene-to-trait associations in the Zhang study and this study, where we select 47 random genes 10,000 times.

**Figure S6.**
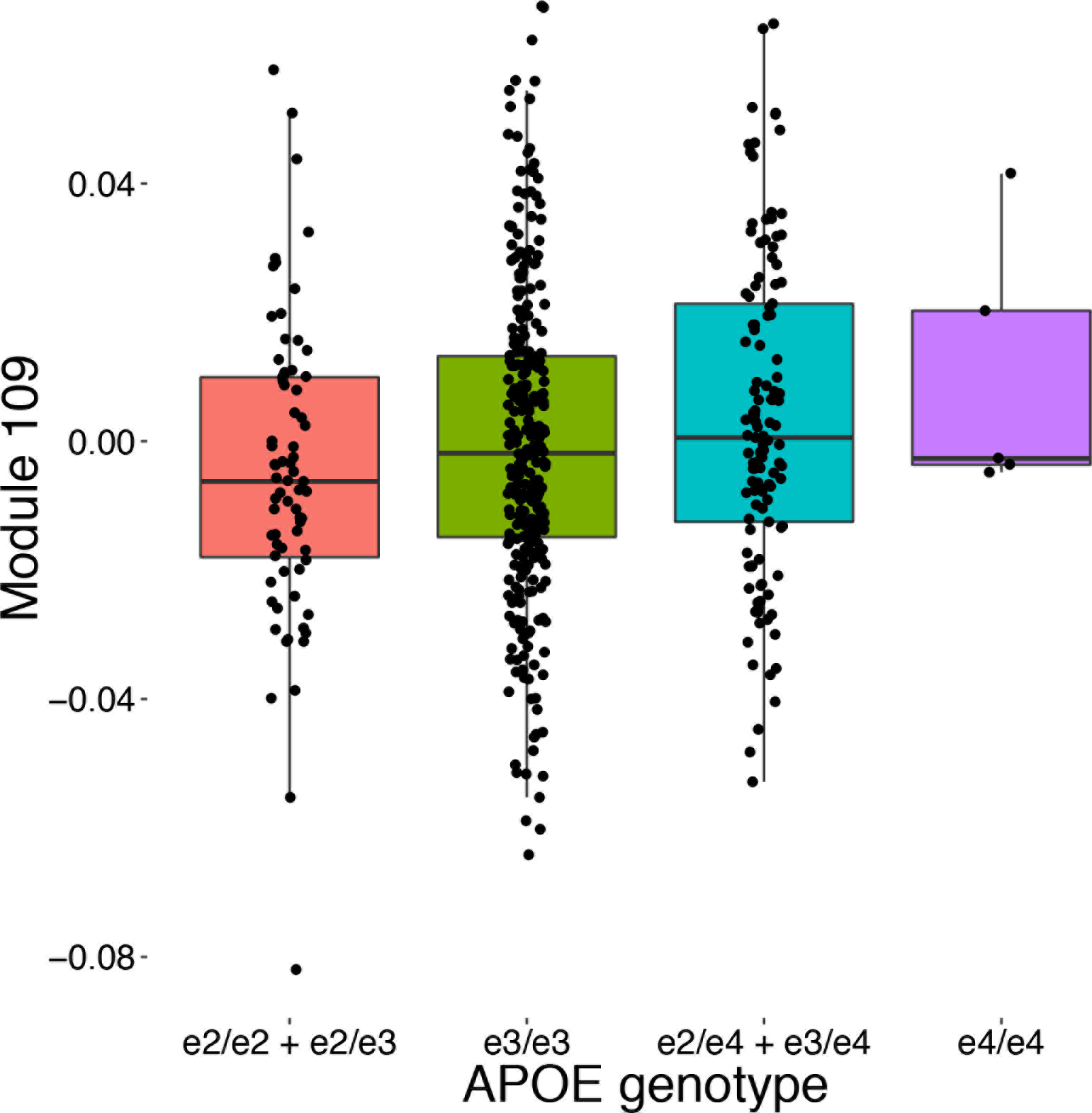
Expression of module 109 meta-feature categorized in order of AD risk, by APOE genotype status. Due to small sample sizes, genotypes e2/e2 and e2/e4 were collapsed into larger categories. APOE genotype status is defined as: e2 (rs7412-T, rs429358-T), e3 (rs7412-C, rs429358-T), e4 (rs7412-C, rs429358-C). Each dot represents one subject.

